# Frequency shifts and depth dependence of premotor beta band activity during perceptual decision-making

**DOI:** 10.1101/306753

**Authors:** Chandramouli Chandrasekaran, Iliana E. Bray, Krishna V. Shenoy

**Author notes:** both authors contributed equally to this work (Chandramouli Chandrasekaran, Iliana E. Bray).

## Abstract

Neural activity in the premotor and motor cortex shows prominent structure in the beta frequency range (13-30 Hz). Currently, the behavioral relevance of beta band activity (BBA) in premotor and motor regions is not well understood. The underlying source of motor BBA and how it changes as a function of cortical depth is also unknown. Here, we addressed these unresolved questions by investigating BBA recorded using laminar electrodes in the dorsal premotor cortex (PMd) of two male rhesus macaques performing a visual reaction time (RT) reach discrimination task. We observed robust BBA before and after the onset of the visual stimulus but not during the arm movement. While post-stimulus BBA was positively correlated with RT throughout the beta frequency range, pre-stimulus correlation varied by frequency. Low beta frequencies (~15 to 20 Hz) were positively correlated with RT and high beta frequencies (~25 to 30 Hz) were negatively correlated with RT. Simulations suggested that these frequency-dependent correlations could be due to a shift in the component frequencies of the pre-stimulus BBA as a function of RT, such that faster RTs are accompanied by greater power in high beta frequencies. We also observed a laminar dependence of BBA, with deeper electrodes demonstrating stronger power in low beta frequencies both pre- and post-stimulus. The heterogeneous nature of BBA and the changing relationship between BBA and RT in different task epochs may be a sign of the differential network dynamics involved in expectation, decision-making, and motor preparation.

## SIGNIFICANCE STATEMENT

Beta band activity (BBA) has been implicated in motor tasks, in disease states, and as a signal for brain-machine interfaces. However, the functional role of BBA and its laminar organization in motor cortex are poorly understood. Here we addressed these unresolved issues using simultaneous recordings from multiple cortical layers of the motor cortex of monkeys performing a decision-making task. Our key finding is that BBA is not a monolithic signal. Instead, BBA seems to be composed of at least two frequency bands. The relationship between BBA and eventual behavior, such as reaction time, also dynamically changes depending on task epoch. We also found that BBA is laminarly organized, with greater power in deeper electrodes for low beta frequencies.

## INTRODUCTION

Fluctuations in the beta (13-35 Hz) range of the local field potential (LFP) and spiking activity are consistently observed in monkeys performing instructed delay (Sanes and Donoghue, 1993; Zhang et al., 2008; Kilavik et al., 2012, 2013; Stetson and Andersen, 2014; Khanna and Carmena, 2017) and cognitive tasks (Murthy and Fetz, 1992; Lee, 2003; Buschman and Miller, 2007; Pesaran et al., 2008; DePasquale and Graybiel, 2015; Sherman et al., 2016; Haegens et al., 2017). Other studies demonstrated prominent BBA in humans performing motor and cognitive tasks (Rubino et al., 2006; Tzagarakis et al., 2010; Zaepffel et al., 2013). Clinical studies suggest that BBA changes with age (Rossiter et al., 2014b), is modulated in disease states (Brown, 2006; Brittain et al., 2014; Rossiter et al., 2014a; Proudfoot et al., 2017), and may be useful for brain machine interfaces (Bai et al., 2008; Flint et al., 2013; So et al., 2014; Gilja et al., 2015; Stavisky et al., 2015; Pandarinath et al., 2017a). Despite insights gained about BBA, questions about its role and origin still remain. Here, we focus on two unresolved questions.

First, we wanted to understand the relevance of BBA in the motor system for decisionmaking. Three hypotheses have been proposed for the role of BBA in the motor system – postural holding, maintenance of the current state, and attention – each making specific predictions relating BBA and RT (Figure 1, Khanna and Carmena, 2015). The postural holding hypothesis posits that BBA is related to keeping the hand still during the hold period of instructed delay tasks (Baker et al., 1999; Kristeva et al., 2007). A second hypothesis suggests that BBA represents the desire to maintain the current state of being (e.g., resisting start of movement) (Gilbertson et al., 2005; Pogosyan et al., 2009; Engel and Fries, 2010). The attentional hypothesis emerged from the study of reach-target selection tasks and suggests that BBA reflects attention (Bouyer et al., 1987; Murthy and Fetz, 1992; Zhang et al., 2008; Saleh et al., 2010). Here, we addressed the behavioral relevance of BBA by examining the relationship between RT and BBA recorded from PMd of two monkeys (Zhang et al., 2008; Saleh et al., 2010; Tzagarakis et al., 2010; Kilavik et al., 2012; Khanna and Carmena, 2017). The monkeys performed a visual reach decision-making task that engaged their attention, involved the somatomotor system, and induced significant RT variability beyond the variability induced by the different stimulus difficulties.

**Figure 1-.**
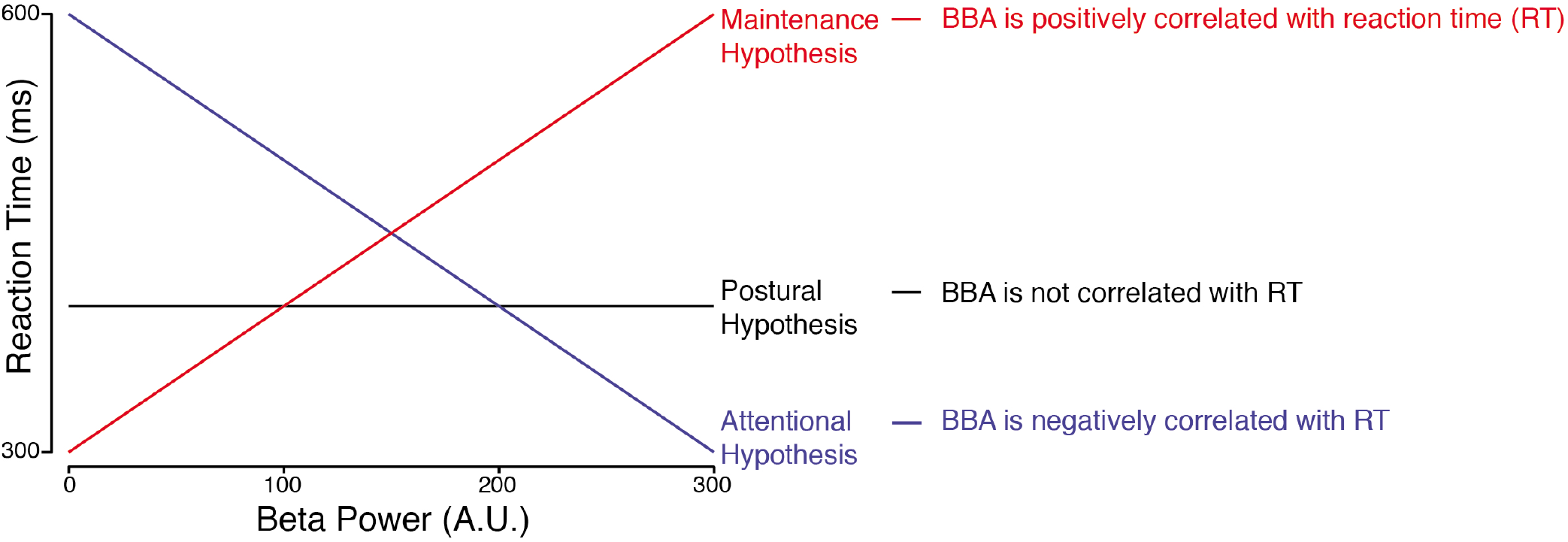
Existing hypotheses about the role of BBA. Three existing hypotheses about the relation between pre-stimulus BBA and RT. The black, horizontal line corresponds to the postural hypothesis of BBA, where there is no relationship between BBA and RT. The red, positively sloped line corresponds to the maintenance hypothesis of BBA, because increased BBA would be tied with longer RTs. The blue, negatively sloped line corresponds to the attentional hypothesis of BBA, because more BBA would be tied to greater attention on the task and therefore shorter RTs. Each dot in the figure is a random, hypothetical RT and beta power used to illustrate the relationship between the two.

Second, we wanted to improve on the currently vague description of the laminar organization of BBA in premotor and motor cortex. Some studies suggest that neurons in deeper cortical layers of M1 (especially layer V) are involved in the generation of BBA (Wetmore and Baker, 2004; Chen and Fetz, 2005; Roopun et al., 2006; Yamawaki et al., 2008). Others suggest that all cortical layers in M1 are involved in BBA (Kondabolu et al., 2016; Sherman et al., 2016). Identifying how BBA changes as a function of cortical depth is needed for developing the next generation of computational models (Kopell et al., 2011; Lee et al., 2013; Bhatt et al., 2016; Sherman et al., 2016). To study the laminar organization of BBA, we used multi-contact electrodes that provided simultaneous recordings across different cortical depths.

We observed that both pre- and post-stimulus BBA was correlated to RT, thus ruling out the postural holding hypothesis. Post-stimulus BBA was positively correlated with RT throughout the 13-35 Hz range, while the correlation between RT and pre-stimulus BBA was positive in the low beta frequencies (∼15 to 20 Hz) and negative in the high beta frequencies (∼25 to 35 Hz). Through simulation, we identified that frequency-dependent correlations between RT and prestimulus LFP power spectra could potentially emerge from a shift in pre-stimulus BBA to higher frequencies for faster RTs. We also found that power spectra of LFPs recorded in deeper electrodes demonstrated more power in low beta frequencies both pre- and post-stimulus.

## METHODS EXPERIMENTAL DESIGN

Here we provide a brief description of the experimental design. Additional details about training and surgery in addition to a description of single neuron responses during the various epochs are found in a previous study (Chandrasekaran et al., 2017). This study focuses on analysis of the pre-stimulus and post-stimulus LFP recorded during the same experiments.

### Subjects

Our experiments were conducted using two adult male macaque monkeys (Macaca mulatta; Monkey T, seven years, 14 kg; O, eleven years, 15.5 kg) trained to reach for visual targets for a juice reward. Monkeys were housed in a social vivarium with a normal day/night cycle. The protocols for our experiments were approved by the Stanford University Institutional Animal Care and Use Committee. We initially trained monkeys to come out of the cage and sit comfortably in a chair. After initial training, we performed sterile surgeries during which monkeys were implanted with head restraint holders (Crist Instruments, cylindrical head holder) and standard recording cylinders (Crist Instruments). Cylinders were centered over caudal PMd (+ 16, 15 stereotaxic coordinates) and placed surface normal to the cortex. We covered the skull within the cylinder with a thin layer of dental acrylic/palacos.

### Apparatus

The general set-up for the experiments is shown in Fig. 2a. Monkeys sat in a customized chair (Crist Instruments, Snyder Chair) with the head restrained via the surgical implant. The arm not used for reaching was gently restrained using a tube and a cloth sling. Experiments were controlled and data collected under a custom computer control system (xPC target and Psychophysics Toolbox). Stimuli were displayed on an Acer HN2741 computer screen placed approximately 30 cm from the monkey. A photodetector (Thorlabs PD360A) was used to record the onset of the visual stimulus at a 1 ms resolution. Each session we taped a small reflective hemispheral bead (11.5 mm, NDI Digital passive spheres) to the middle digit of the right hand (left hand for Monkey O). The bead was taped 1 cm from the tips of the fingers, and the position of this bead was tracked optically in the infrared (60 Hz, 0.35 mm root mean square accuracy; Polaris system; Northern Digital). Eye position was tracked with an overhead infrared camera (estimated accuracy of 1°, Iscan, Burlington, MA). To get a stable eye image for the overhead infrared camera which acquires the eye image, an infrared mirror transparent to visible light was positioned at a 45° angle (facing upwards) immediately in front of the nose. This mirror reflected the image of the eye in the infrared range while letting visible light pass through. A visor placed around the chair prevented the monkey from bringing the bead to his mouth or touching the infrared mirror or the juice tube.

**Figure 2-.**
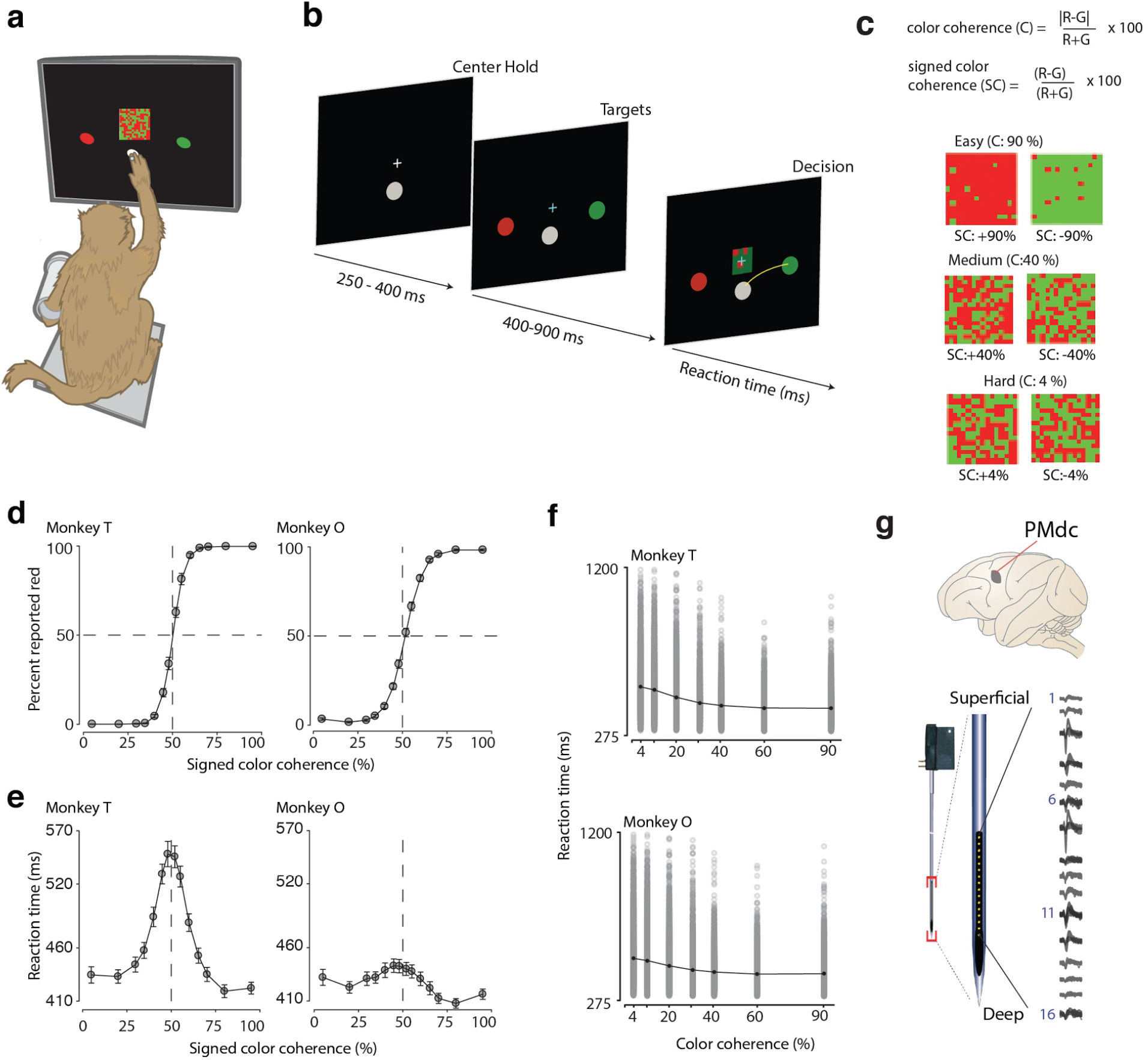
Recording locations, techniques, task, and discrimination behavior. **a**: An illustration of the experimental setup for data gathering in the discrimination task. We gently restrained the resting arm with a plastic tube and cloth sling. We tracked a reflective IR bead taped on the middle digit of the unrestrained hand to mimic a touch screen and to provide an estimate of instantaneous arm position. We tracked eye position using an infrared reflective mirror placed in front of the monkey’s nose. **b**: Example timeline of the discrimination task. **c**: Examples of different stimulus ambiguities used in the experiment parameterized by the color coherence of the checkerboard defined as 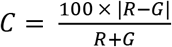. The corresponding signed color coherence is defined as 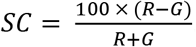. Positive values of signed color coherence denote more red than green squares and vice-versa. **d-e**: Average discrimination performance (d) and reaction time (RT) (e) over sessions of the two monkeys as a function of the signed color coherence of the checkerboard. RT plotted here includes both correct and incorrect trials for each session and then averaged across sessions. Gray markers show measured data points along with 2x(standard error) estimated over sessions, though variation is so small that they are difficult to see in (d). The black line segments are drawn in between these measured data points to guide the eye. For most data points in (d), the error bars lie within the markers. X-axes in both (d) and (e) depict the signed color coherence in %. Y-axes depict the percent responded red in (d) and RT in (e). Also shown in (d) are discrimination thresholds (M±SD over sessions) estimated from a Weibull fit to the overall percent correct as a function of coherence. The discrimination threshold is the color coherence level at which the monkey made 81.6% correct choices. 24 sessions for monkey T (47483 trials) and 44 sessions for monkey O (70,250 trials) went into these averages. **f**: RT as a function of checkerboard coherence. For each coherence, the mean RT is shown in black and connected linearly, with gray markers showing individual RTs. There is large variation of RTs both across and within coherences. **g**: Location of PMd along with an example recording from a 16 electrode, 150 μm spacing U-probe.

### Task structure

Experiments consisted of a sequence of trials, which each lasted a few seconds; successful trials resulted in a juice reward, and unsuccessful trials resulted in a timeout lasting 2-4 seconds. An example trial timeline is shown in Fig. 2b. Monkeys used their unrestrained arm (Monkey T used his right arm, Monkey O used his left arm) to reach to touch either red or green targets based on the dominant color in a central, static checkerboard cue composed of isoluminant red and green squares. For every trial, the monkey placed his unrestrained arm on a central target (diameter = 24 mm) and fixated on a small white cross (diameter = 6 mm). After ∼350-400 ms had elapsed, two isoluminant colored targets appeared 100 mm to the right and left of the central target. The target configuration was randomized so that colors were not always tied to reach directions: sometimes the red target was on the left and green on the right, while other trials had the opposite configuration. After an additional hold period (varying from 400 to 900 ms), a static checkerboard cue (15 x 15 grid of squares; each square 2.5 mm x 2.5 mm) composed of isoluminant red and green squares appeared on the screen around the fixation cross (example stimuli are shown in Fig. 2c). The monkeys reached for the target whose color matched the dominant color in the central checkerboard cue. For example, when there was more green than red in the central checkerboard cue, the monkey had to choose the green target. To “choose” a target, the animals moved their hand from the central hold point and stably held a target for a short duration (minimum of 200 ms). The task was an RT paradigm, so the monkeys were free to initiate their reach whenever they felt there was sufficient evidence for them to provide a response. We did not impose any delayed feedback procedure in this task such as a delay between the time of reward and the completion of a reach for a correct target. The juice reward was provided to the monkey immediately after the monkey provided a correct response (Roitman and Shadlen, 2002).

We parameterized the checkerboard cue at several different levels from almost fully red to almost fully green. We used 14 levels of red (ranging from 11 red squares to 214 red squares) in the central checkerboard cue. Each level of red had a complementary green level (e.g., 214 R, 11 G; and 214 G, 11 R-squares). This defined seven levels of color coherence

(defined as 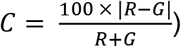), ranging from 4 – 90%. The corresponding signed color coherence was estimated without taking the absolute value of the difference 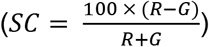. For Monkey T, we used a uniform distribution of hold period durations between the onset of the targets and the onset of the checkerboard cue. Monkey O attempted to anticipate the checkerboard cue onset. To minimize this anticipation and reduce predictability we used an exponential hold period duration (400 – 800 ms) between the onset of the targets and the onset of the checkerboard cue.

### Electrophysiological Recordings

Stereotactic coordinates, known response properties of PMd and M1, and neural responses to muscle palpation served as our guides for electrophysiological recordings. We placed the chambers surface normal to the cortex to align with the skull of the monkey, and recordings were performed perpendicular to the surface of the brain. Recordings were made anterior to the central sulcus, lateral to the spur of the arcuate sulcus, and lateral to the precentral dimple. For both monkeys, we confirmed our estimate of the upper and lower arm representation by repeated palpation at a large number of sites to identify muscle groups associated with the sites. Monkey T used his right arm to perform tasks while O used his left arm. Recordings were performed in PMd and M1 contralateral to the arm used by the monkey.

We performed linear multi-contact electrode (U-probe) recordings in the same manner as single electrode recordings with some minor modifications. We used a slightly sharpened guide tube to allow the U-probe to penetrate the Dura more easily. We also periodically scraped away any overlying tissue on the dura under anesthesia. Sharp guide tubes and scraping away dura greatly facilitated penetration of the U-probe. We typically penetrated the brain at very slow rates (∼2 – 5 μm/s). Once we felt that we had a reasonable sample population of neurons potentially spanning different cortical layers, we stopped and waited for 45-60 min for the neuronal responses to stabilize. The experiments then progressed as usual. We used 180 μm thick, 16-electrode U-probes with an inter-electrode spacing of 150 μm; electrode contacts were ∼100 kΩ in impedance.

We attempted to minimize the variability in U-probe placement on a session-by-session basis so that we could average across sessions. Our approach was to place the U-probe so that the most superficial electrodes (electrodes 1, 2 on the 16 channel probe) were able to record multi-unit spiking activity. Any further movement of the electrode upwards resulted in the spiking activity for those electrodes disappearing and a change in the overall activity pattern of the electrode (suppression of overall LFP amplitudes). Similarly, driving the electrodes deeper resulted in multiphasic extracellular waveforms and also a change in auditory markers which were characterized by decreases in overall signal intensity and frequency content; both markers suggested that the electrode entered white matter (Cooper et al., 1969). We utilized these physiological markers as a guide to place electrodes and thereby minimize variability in electrode placement on a session-by-session basis. Recording yields and this careful electrode placement were in general better in monkey T (average of ∼16 units per session) than monkey O (average of ∼9 units per session). Random placement of U-probes on a day-to-day basis would flatten out the average visuomotor index and dilute or entirely remove significant differences in the discrimination time differences between superficial and deep electrodes.

The insertion technique necessitated a careful watch over the electrode while lowering to ensure that it did not bend, break at the tip or excessively dimple the dura. We therefore were unable to use a grid system to precisely localize the location of the U-probes on different days and to provide a map of how laminar profiles change in the rostrocaudal direction.

### Local field potentials

LFP recordings in T were performed using a 2 KHz sampled signal. We then resampled this signal at 1 KHz and performed subsequent spectral analysis on appropriate time epochs. For monkey O, two methods were used. For 17 of the sessions, we recorded LFP at 2 KHz, as in T. For the remaining 27 sessions, we recorded broadband extracellular activity at 30 KHz. We resampled this broadband extracellular signal at 1 KHz and then again used it for subsequent spectral analysis. All resampling was performed using the MATLAB resample command that first applies a delay compensating low pass filter and then subsequently resamples the data avoiding antialiasing.

### Reaction Time

Reaction time (RT) is defined as the time between stimulus onset and the monkey’s selection of a target. RT is described in units of milliseconds. A reaction time less than or equal to 300 ms indicates that the monkey did not incorporate the presented stimulus into his response. These trials are not representative of decision-making based on the provided stimulus and were therefore removed from our analysis.

## STATISTICAL ANALYSIS

### Psychometric curves for accuracy

For the analysis of the behavior, we used the same 24 sessions for monkey T (47,483 trials) and 44 sessions for monkey O (70,250 trials) from which we examined electrophysiological data. Fits to psychometric curves and RT regressions were performed on a per-session basis and then averaged over sessions. The behavior of an average session was estimated from ∼1500 trials. RT was estimated for each session by including both correct and incorrect trials for each signed color coherence.

We fit psychometric curves that describe how discrimination accuracy changed as a function of color coherence. For every experiment, we estimated the monkey’s sensitivity to the checkerboard cue by estimating the probability (*p*) of a correct choice as a function of the color coherence of the checkerboard cue (*C*). We used the psignifit toolbox to fit this accuracy function using a Weibull cumulative distribution function (Wichmann and Hill, 2001):

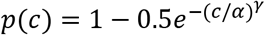

The discrimination threshold, *α*, is the color coherence level at which the monkey would make 81.6% correct choices. The second parameter, *γ*, describes the slope of the psychometric function. The mean*α*parameter across sessions was used as the threshold. We fit threshold and slope parameters on a session-by-session basis and averaged the estimates. The mean and standard deviation of the threshold estimates are reported in Fig. 2d.

### RT vs. coherence

To examine if RT changed with color coherence, we adopted the procedure from (Roitman and Shadlen, 2002) and used a linear regression between RT and log coherence.

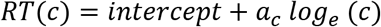

We fit this regression model (Fig. 2e) with *a_c_* as the slope of the regression.

### Power spectra

To estimate the power spectra, we used the Chronux toolbox for MATLAB (Mitra and Bokil, 2008; Mitra et al., 2016) which implements the multi-taper spectral estimation method, with a time-bandwidth product of three and with five leading tapers. Choice of other tapers did not result in any changes in our conclusions. We removed the DC offset from the LFP time series and used a second-order 11R notch filter to remove line noise (Mitra and Bokil, 2008; Mitra et al., 2016). Line noise, which is centered at 60 Hz, arises from radiative electrical pickup from lights and power sockets, currents due to ground loops, and currents induced by magnets in DC power supplies in the experimental setup (Mitra and Bokil, 2008). We centered the filter at 60 Hz and set the quality factor (related to the filter bandwidth) to 35. The power spectra have arbitrary units (A.U.) before they are normalized.

We only plot the power spectra from 2 Hz to 50 Hz. We saw no significant activity in the range of 50 Hz to 500 Hz. For the normalized power spectra from 2 to 90 Hz, the Z scores from 50 Hz through 90 Hz were below zero for all analyzed periods of the task (pre-stimulus, poststimulus, and post-movement).

### Normalization of power spectra

For each trial, we normalized the power spectrum over all power values (for each frequency for all electrodes) from all trials in that session. We calculated the Z Score by subtracting the mean (of all power values from all trials in that session) from each point and dividing by the standard deviation.

### Standard Error

Standard error was defined as 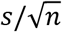, where *s* is the standard deviation of the power spectra for several sessions with respect to the sessions, and *n* is the number of sessions. Standard error is shown in shading in plots of power spectra.

### Split into RT quantiles

We first calculated the breaks for the RT percentiles for that monkey on that session, separating the trials with RTs either greater than 85% of trials in that session and the trials with RTs smaller than 15% of trials in that session. We then averaged the normalized power spectra over trials within each RT quantile and finally averaged over all electrodes within each quantile. Then within each quantile, we averaged over all sessions, giving two normalized grand average power spectra each over all trials, channels, and sessions.

### Correlation between BBA and RT

For each electrode, per session of data (several trials), we computed the partial Spearman correlation between the normalized power at each frequency with reaction time, controlling for the coherence of the checkerboard. We then averaged the correlations over all electrodes and all sessions. Significance of the correlation values were adjusted using the Benjamini & Hochberg (Benjamini and Hochberg, 1995) procedure for controlling the false discovery rate (FDR) of a family of hypothesis tests (Groppe, 2016).

We decided to do a partial correlation in order to control for the confounding variable, the coherence of the checkerboard, which we know affects the RT and also likely affects the LFP power spectra and would therefore have otherwise given misleading correlation values.

### Simulating Relationships between BBA and RT

In order to clarify the mathematical relationship between BBA and RT, we ran a series of simulations (Fig. 6). We first randomly generated an RT value within the range typically observed for our monkeys. Then, we created a variety of LFP signals in which the frequency and amplitude were either constant or related in some way to the RT that was generated. The relationship between frequency, amplitude, and RT are specified in the equations below, where randn signifies a random number drawn from the normal distribution. Within each frequency and amplitude relationship, we generated one thousand RTs and corresponding LFP signals. We then calculated the power spectrum for each simulated LFP signal before correlating the power spectra to the randomly generated RT. Each frequency and amplitude relationship resulted in a different correlation with RT. The equations below match the panels shown in Fig. 6.

i. Frequency = 28 + 1.2*randn + .003*RT; Amplitude = 1;
ii. Frequency = 28 + 1.2*randn – .003*RT; Amplitude = 1;
iii. Frequency = 28 + 1.2*randn – .003*RT; Amplitude = .3 + 5e-6*RT;
iv. Frequency = 28 + 1.2*randn – .003*RT; Amplitude = .3 – 5e-6*RT;
v. Frequency = 28 + 1.2*randn; Amplitude = .3 + 5e-6*RT;
vi. Frequency = 28 + 1.2*randn; Amplitude = .3 – 5e-6*RT;

## RESULTS

Two trained monkeys (T and O) discriminated the dominant color of a central, static checkerboard cue composed of mixtures of red and green squares and used an arm movement to report the decision (Fig. 2a, Coallier et al., 2015). Fig. 2b depicts a trial timeline. The trial began when the monkey touched the center target and fixated on the cross. After a variable target viewing period, the red-green checkerboard cue appeared. The task of the monkey was to make an arm movement toward the target (red vs. green) that matched the dominant color of the checkerboard cue. We parameterized difficulty of the discrimination (example stimuli shown in Fig. 2c) by a color coherence measure (C) defined as the absolute difference in the number of red and green squares normalized by the total number of squares in the checkerboard (C = 100*|R-G|/(R+G)). A corresponding signed color coherence measure (SC) is defined as SC = 100*(R-G)/(R+G). We previously reported the behavior of the monkeys while they performed this task (Chandrasekaran et al., 2017). Here we present the psychometric and chronometric curves for the sessions where we examined the LFP.

On average across sessions, decreases in color coherence resulted in more errors (Fig. 2d). We fit the proportion correct as a function of unsigned coherence (C) using a weibull function to estimate slopes and thresholds (average R^2^, T: .99 (over 24 sessions, 47483 trials), O: .99 (over 44 sessions, 70250 trials), slope (β), M±SD over sessions, T: 1.30± 0.16, O: 1.26± 0.15). Monkey T displayed more sensitivity than Monkey O (thresholds are computed on a per-session basis and averaged over sessions at 81.6% correct, (M±SD): T, 9.87%±1.12%, O: 15.05±1.79%, two-tailed test, Wilcoxon rank sum comparing median thresholds, p=1.292e-11).

A decrease in color coherence also resulted in a slower mean RT (Fig. 2e, using a regression to test if mean RT increases as log_e_ coherence decreases (harder stimulus difficulties as in (Roitman and Shadlen, 2002); average R^2^, T: 0.94, O: 0.59; slope of regression: M±SD over sessions, T: −41.1±6.3 ms/log_e_ coherence (%), O: −8.6±4.5 ms/log_e_ coherence (%)). Monkey T had a larger range of RTs compared to Monkey O (Comparing the RT range between easiest and hardest difficulties (M±SD) estimated over sessions; T: 115±19 ms and O: 28±11 ms, Wilcoxon ranksum comparing median ranges of RT, p=1.292e-11).

Although color coherence explains considerable variation in RT, there is significant variation that is not explained by the coherence. A linear regression between RT and stimulus coherence only explained 10.8% of the variance in monkey T and only 1.3% in monkey O. Variation in RT is readily apparent even within a given color coherence (Fig. 2f). Our hypothesis is that this RT variability is at least in part related to fluctuations in BBA (See Figure 1, Pogosyan et al., 2009; Kilavik et al., 2012; Khanna and Carmena, 2017).

### LFP and neuronal responses during the pre-stimulus period show prominent beta band activity

We first examined our LFPs recorded in PMd, specifically examining how the power across different frequencies of the LFP changed throughout the reach decision task. BBA is apparent in the pre-stimulus period (600 ms before the appearance of the checkerboard the stimulus), decreases during the decision-formation period, and remains low during the movement epoch (Fig. 3a, 3b). This pre-stimulus increase in power in the 15-35 Hz range is consistent with the definition of BBA in both frequency (from 15 to 35 Hz) and timing within task behavior (Sanes and Donoghue, 1993; Baker et al., 1997; Kilner et al., 1999; Riddle and Baker, 2006; Rubino et al., 2006; Baker, 2007; Klostermann et al., 2007; Chakarov et al., 2009; Zaepffel et al., 2013). Decreases in BBA after movement onset are also consistent with these and other prior reports of beta event related desynchronization. Finally, activity in the delta band (0.5 to 4 Hz), theta band (4 to 7 Hz), and alpha band (8 to 12 Hz) are present both before and after checkerboard onset (Fig. 3a). We found that there was essentially no activity in the gamma band (40-100 Hz) (Fig. 3a).

**Figure 3-.**
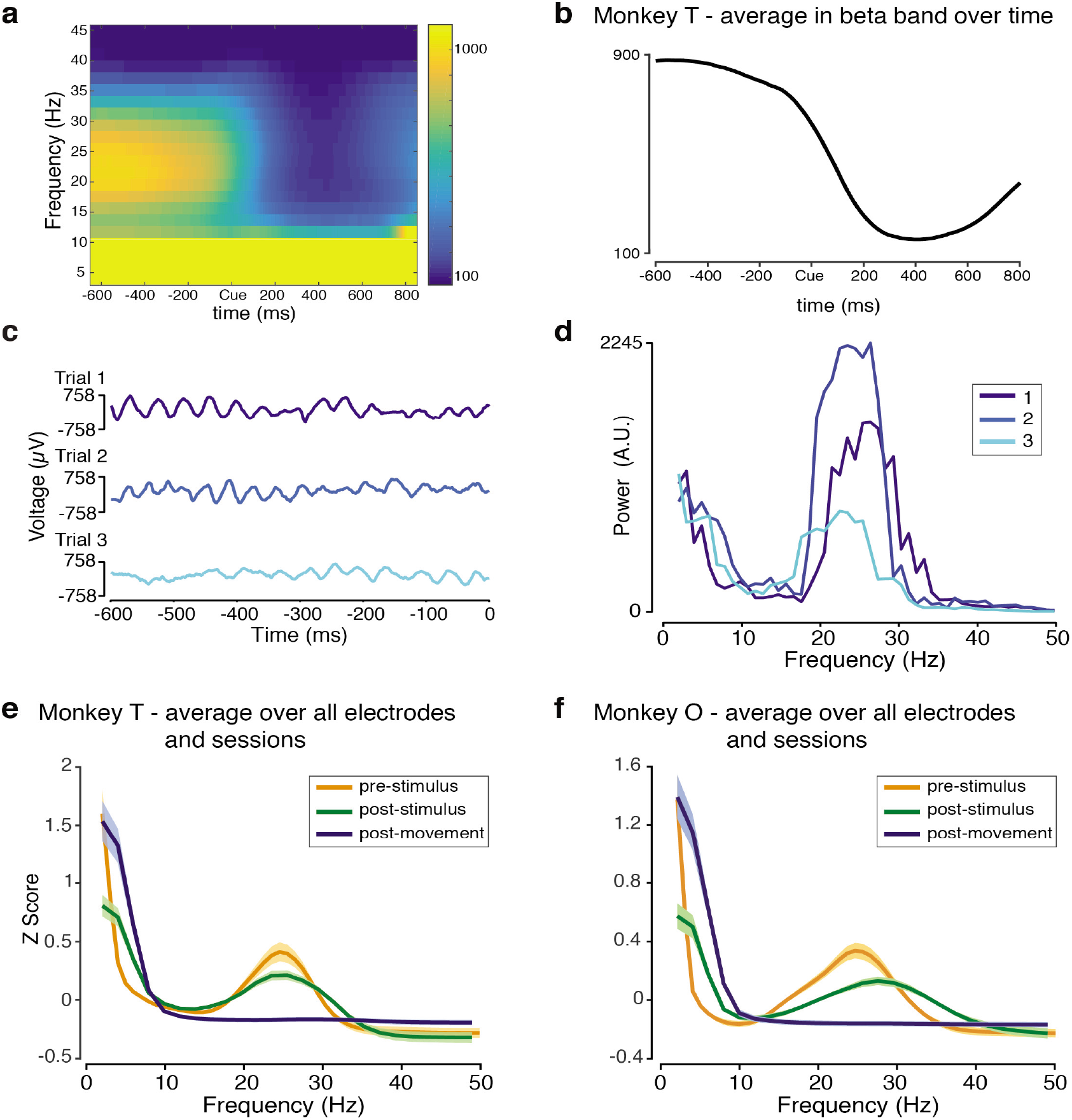
Existence of BBA during hold period before the visual stimulus. **a**: Spectrogram aligned to checkerboard onset (indicated with Cue), averaged over all electrodes, trials, and sessions for Monkey T. The Y-Axis represents frequency and is shown in Hertz. The X-axis represents time in milliseconds. Color represents power in arbitrary units (A.U.). Clear presence of pre-stimulus BBA is seen, with lower-power post-stimulus BBA. **b**: Activity in the beta band (13-30 Hz) over time, averaged over all electrodes, trials, and sessions for Monkey T. The Y-Axis is power in (A.U.) and the X-axis represents time in milliseconds. **c**: The LFP time series of three trials of Electrode 2 during a single session. The colors are unique to each trial and consistent with subplot (d). The time series are shown as microvolts per millisecond. **d**: Power spectra of three example trials during the epoch before the checkerboard. Power in (A.U.) is plotted against frequency (Hertz). **e-f**: Normalized power spectra of the LFP during the epoch before the checkerboard (orange), after the checkerboard (green), and after movement (blue). (e) Monkey T grand average over all electrodes, trials and sessions. (f) Monkey O grand average over all electrodes, trials, and sessions. The power spectra have been normalized, and their Z Scores are plotted against frequency (Hertz). Standard error over sessions is shaded.

Several other analyses confirmed the existence of BBA during the pre-stimulus period. Temporal fluctuations in the beta band were readily visible in individual trials of the LFP suggesting that we are not artificially separating a broadband signal into signals of a specific frequency (Fig. 3c). The power spectra for the trials shown in Fig. 3c corroborated this observation of signals in the 15-35 Hz range (Fig. 3d). Finally, pre-stimulus BBA was consistently observed in our population recordings (Fig. 3e & 3f). Figs. 3e & 3f plot the average power spectrum over all trials, electrodes, and sessions for three different task periods: pre-checkerboard cue, post-checkerboard cue, and post-movement. Both monkeys show significant BBA during the pre-stimulus period, each with peak frequencies slightly below 30 Hz.

Across both monkeys, BBA observed after the checkerboard (during the post-checkerboard period) differs from pre-stimulus BBA (Fig. 3e & 3f). After the checkerboard, BBA has decreased peak power and a broader peak (covering more frequencies). The frequencies present are still consistent with the frequency definition of BBA.

### RT covaries with BBA frequency and power

Our first goal in this study was to better understand the relationship between BBA from the pre- and post-stimulus periods and behavior. First, we examined if there were significant relationships between pre-stimulus BBA and RT. As an initial, exploratory analysis, we examined the extremes of the data by splitting the data into the (fastest) trials with the smallest 15% of RTs and the (slowest) trials with the largest 15% of RTs and compared the average power spectra of the two groups for each monkey. Using the 5th and 95th percentiles suggested similar patterns. Across both monkeys during the pre-stimulus period, we found that the faster RTs have more power in the higher frequencies of BBA (approximately 25 to 30 Hz) (Fig. 4a & 4b). In Monkey T, in the lower frequencies of BBA (approximately 15 to 25 Hz), the slower RTs have more power. Combined, this leads to a frequency shift between the RT quantiles, with the power spectra for the slower RT trials slightly shifted towards the lower frequencies. In Monkey O, however, the faster (smallest) RTs have more power for both the low and high frequencies of BBA, so the perceived shift is not present.

To more rigorously quantify this relationship between RT and pre-stimulus BBA, we examined the correlation between these two variables at each and every frequency. We performed this analysis using partial correlations; i.e., we estimated the correlation between prestimulus BBA and RT while using checkerboard coherence as a covariate. We then averaged the partial correlations over the 16 electrodes. Correlation analyses exploiting the simultaneous nature of our recordings were not notably different from the averaging analysis. So we only report the results obtained from averaging partial correlations over electrodes.

**Figure 4-.**
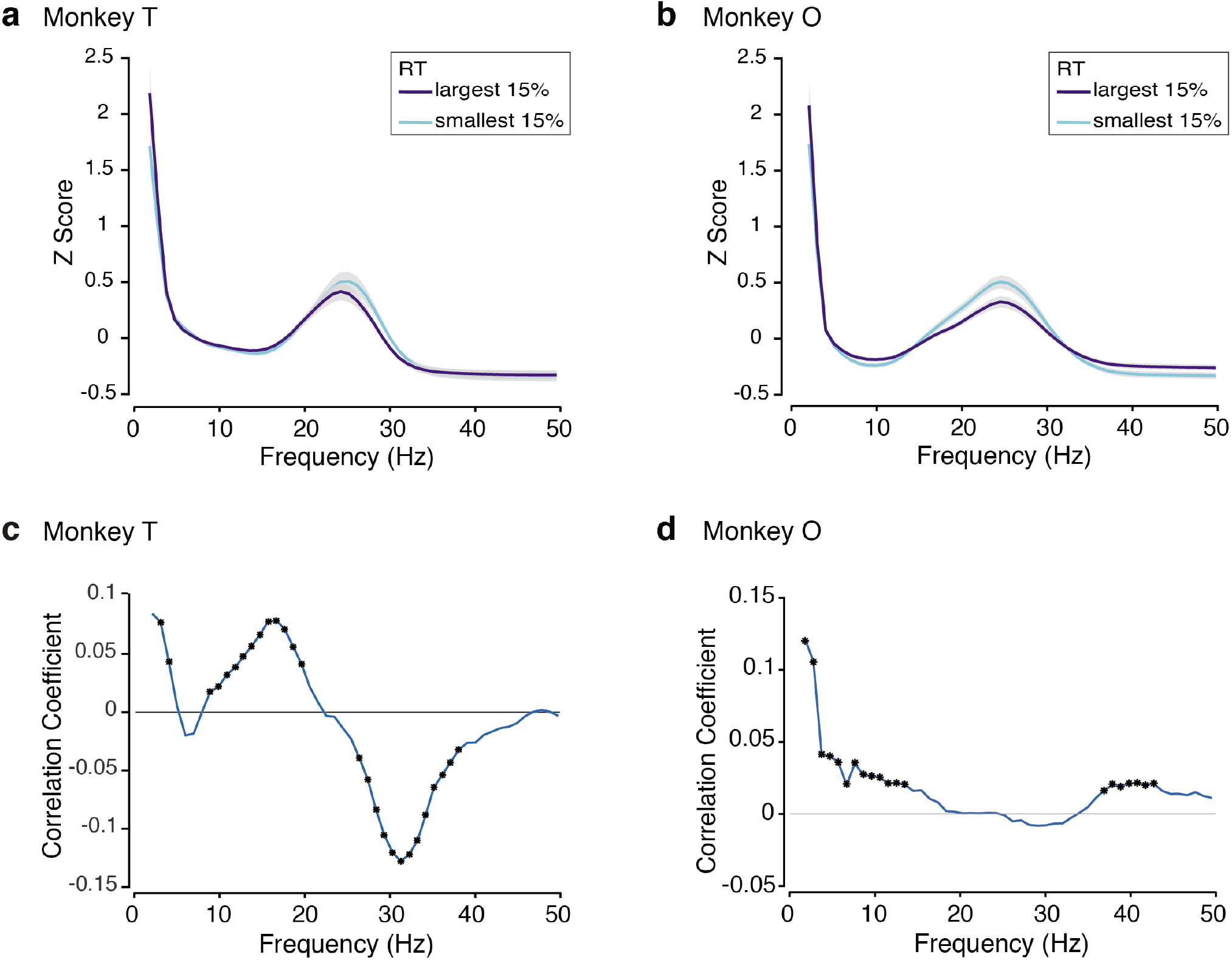
Relation between pre-stimulus BBA and Reaction Time. **a,b**: Normalized pre-stimulus power spectra grouped into two reaction time quantiles and averaged over all trials within that group, all electrodes, and all sessions for Monkey T (a) and Monkey O (b). The two quantiles are the 15% largest (slowest) reaction times and the 15% smallest (fastest) reaction times. The power spectra have been normalized and their Z Scores are plotted against frequency (Hertz). Standard error over sessions is shown in gray. **c,d**: Correlation between normalized pre-stimulus power spectra with RT as a function of frequency for Monkey T (c) and Monkey O (d). Asterisks indicate points along the curve where the correlation is significant (adjusted p-value less than 0.05).

Our analysis suggested a positive correlation between BBA and RT around 15 Hz (T: peak at approx. 16 Hz, r = 0.0785, p = 9.9341e^−7^; O: peak at approx. 12 Hz, r = 0.0214, p = 0.0056) and a negative correlation between BBA and RT around 35 Hz (T: minimum at approx. 31 Hz, r = −0.1278 p = 9.9341e^−7^; O: not significant) (Fig. 4c & 4d). The presence of significant correlations is inconsistent with the postural holding hypothesis. However, varying correlations by frequency support both the maintenance hypothesis (purely positive correlations with BBA) and the attentional hypothesis (purely negative correlations with BBA) within different subregions of the beta band (maintenance for low BBA and attentional for high BBA).

We next performed the same analyses on the post stimulus (post-checkerboard) BBA to better understand its relation to RT. Across both monkeys during the post-stimulus period, we see that the slower (larger) RTs (85th percentile) have more power in the lower frequencies of BBA (approximately 15 to 25 Hz) (Fig. 5a & 5b). In Monkey O, in the higher frequencies of BBA (approximately 25-35 Hz), the faster (smaller) RTs have more power. Combined, this leads to a frequency shift between the RT quantiles, with the power spectra for the slower RT trials slightly shifted towards the lower frequencies. In Monkey T, however, the slower (larger) RTs have more power for both the low and high frequencies of BBA, so the perceived shift is not present. Across both monkeys, the correlation between post-stimulus activity and RT is positive for both low and high beta (as well as some high alpha) (T: peak at approx. 21 Hz, r = 0.13, p = 3.3114e^−7^; O: peak at approx. 21 Hz, r = 0.1167, p = 8.1205e^−13^) (Fig. 5c & 5d). The correlation is negative for gamma activity in the low gamma band (T: minimum at approx. 37 Hz, r = −0.06, p = 3.3114e^−7^; O: minimum at approx. 47 Hz, r = −0.0708, p = 5.8103e^−9^).

**Figure 5-.**
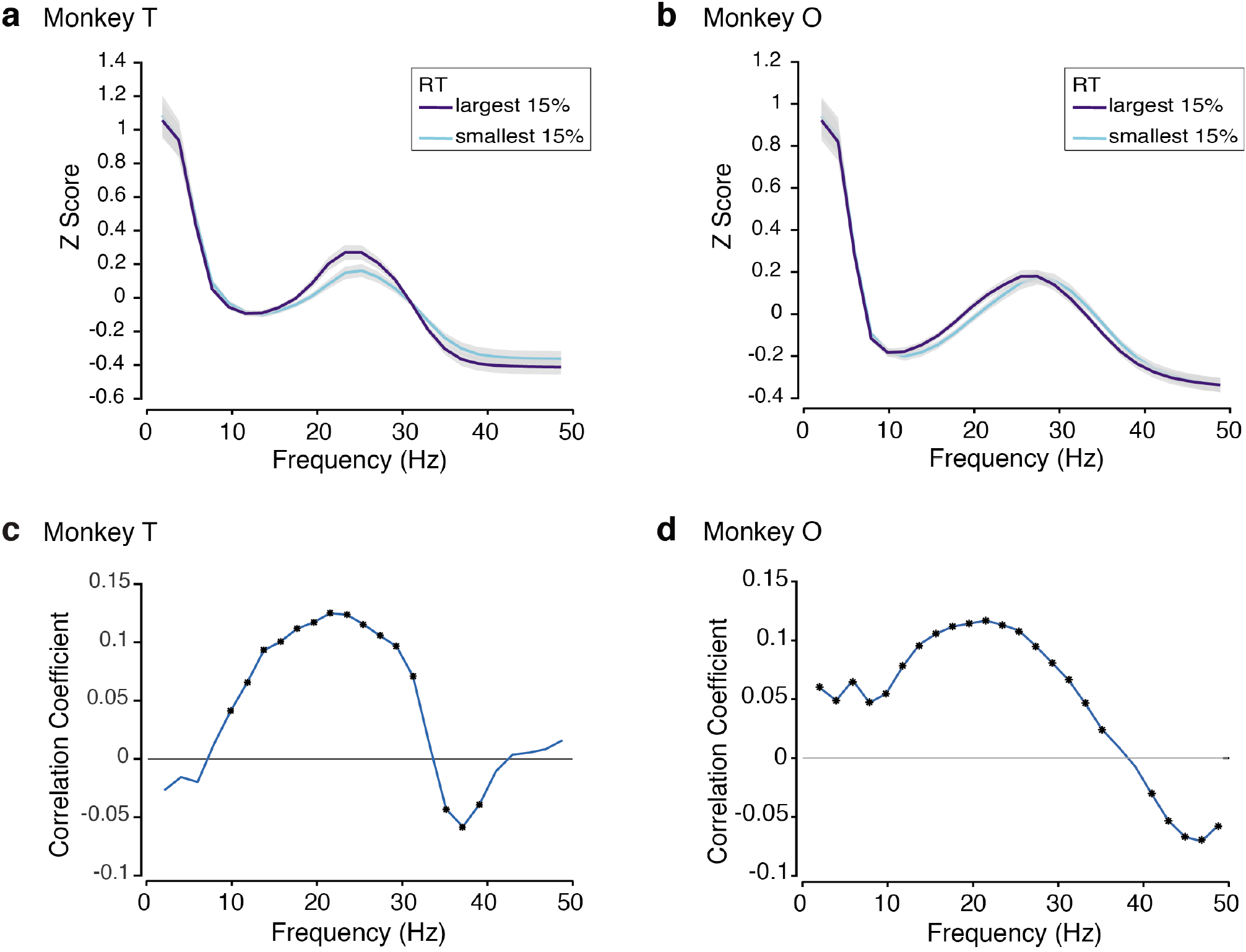
Relation between post-stimulus BBA and Reaction Time. **a,b**: Normalized post-stimulus power spectra grouped into two reaction time quantiles and averaged over all trials within that group, all electrodes, and all sessions for Monkey T (a) and Monkey O (b). The two quantiles are the 15% largest (slowest) reaction times and the 15% smallest (fastest) reaction times. The power spectra have been normalized and their Z Scores are plotted against frequency (Hertz). Standard error over sessions is shown in gray. **c-e**: Correlation between normalized post-stimulus power spectra with RT as a function of frequency for Monkey T (c) and Monkey O (d). Asterisks indicate points along the curve where the correlation is significant (adjusted p-value less than 0.05).

These results for the post-stimulus period can also be more broadly viewed as a shift in the component frequencies of the LFP, this time across multiple frequency bands. That is, on faster RT trials, there is less overall beta band activity and slightly more gamma band activity. The opposite is true for the slower RTs.

### Simulations suggest that a frequency shift in BBA is a plausible mechanism for the observed pattern of correlation

In order to better understand the mechanisms behind the frequency dependent correlation between BBA and RT, we used a simulation analysis. The schematic for this analysis is shown in Fig. 6a. First, we randomly generated RT values within the range of RTs typically observed for our monkeys. Then, based on these values and a variety of governing equations for frequency and amplitude, we simulated LFP signals for these hypothetical trials. The signal was defined as

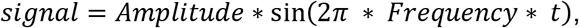

where amplitude and frequency are either constants, linear increasing functions of RT, or linear decreasing functions of RT. We then calculated the power spectra of these simulated signals from these trials and correlated these power spectra to their corresponding RTs. For each group of frequency and amplitude equations, we generated one thousand simulated trials with corresponding RTs, simulated LFP signals, and power spectra. The correlation coefficient as a function of frequency between the simulated power spectra and RTs is shown in Fig. 6b for the six paradigms.

**Figure 6-.**
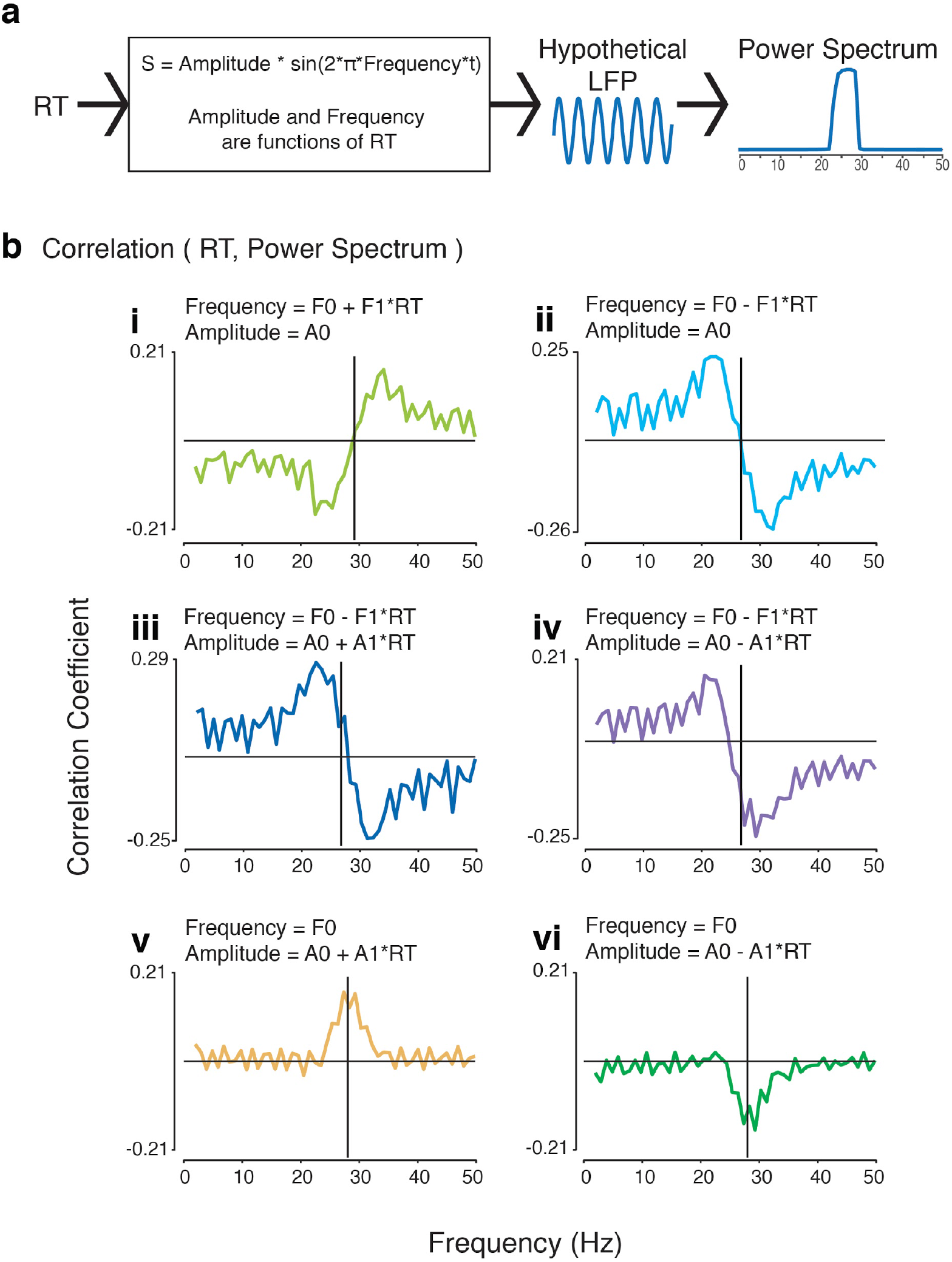
Simulations of Relation between BBA and Reaction Time. Process (a) and results (b) of a simulation that generated synthetic LFP signals as Amplitude*sin(2*pi*Frequency*t). The Amplitude and Frequency of the LFP were defined differently for each case, either as a constant or a function of reaction time. Power spectra were made from these LFP signals, and they were then correlated with RT to create the shown plots of correlation coefficients as a function of frequency for each of the six cases. The amplitude and frequency relationships with RT for each case are shown with the correlations.

The correlations between pre-stimulus BBA and RT observed in the real data (shown in Fig. 4c & 4d) most closely match the correlation when frequency is negatively related to RT. This relationship is robust regardless of the relationship between amplitude and RT (shown in Fig. 6b parts ii, iii, and iv). These findings indicate the presence of a relationship between prestimulus BBA frequency composition and RT, suggesting that pre-stimulus BBA component frequencies are negatively related with RT.

The correlations between post-stimulus BBA and RT observed in the real data (shown in Fig. 5c & 5d) most closely match the correlation when frequency is not related to RT and amplitude is positively related to RT (shown in Fig. 6b, part v). This indicates the presence of a relationship between post-stimulus BBA amplitude and RT with no relationship between poststimulus BBA component frequencies and RT. We do recognize though that additional processes that involve the dynamical balance between beta band activity and gamma band activity can lead to shifts in the frequencies that, in turn, explain the negative correlations in the gamma band but positive correlations in the beta band.

### Deeper cortical layers have stronger activity in the low beta range than the superficial layers

The next goal of our study was to understand how BBA changes as a function of cortical depth. The use of linear multi-contact electrodes (Fig. 2g) provided us with simultaneous recordings across several cortical depths and allowed us to examine whether there was a relationship between cortical depth and BBA.

To examine the degree to which pre-stimulus power in the beta region varied with electrode depth, we divided the electrodes into two groups: the superficial (electrodes 1:8) and the deep (electrodes 9:16). In both monkeys, deeper electrodes (corresponding to deeper cortical layers) have more power around the 10 to 20 Hz region (Fig. 7a & 7b). In one monkey (Monkey O), this pattern of deeper electrodes having more power than surface electrodes continues from approximately 10 Hz until 30 Hz, slightly past its peak frequency (Fig. 7b).

**Figure 7-.**
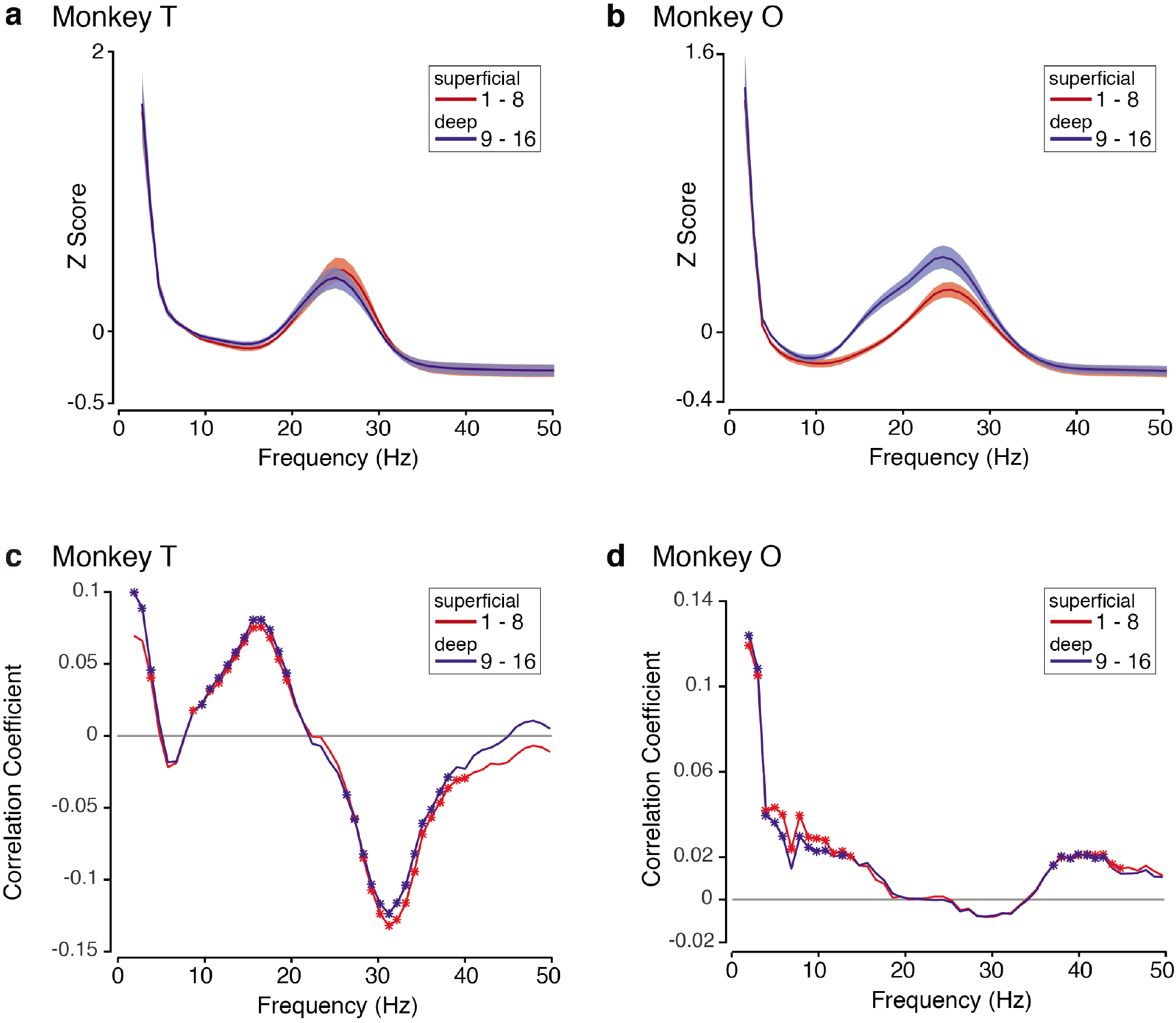
Pre-stimulus BBA by depth. **a,b**: Normalized pre-stimulus power spectra grouped into two electrode groupings and averaged over all trials, all electrodes within that group, and all sessions for Monkey T (a) and Monkey O (b). The power spectra have been normalized, and their Z Scores are plotted against frequency (Hertz). The average over the superficial electrodes is plotted in red, and the average over the deep electrodes is plotted in blue. Standard error over sessions is shaded. **c,d**: Depth dependent correlation between normalized pre-stimulus power spectra with RT as a function of frequency for Monkey T (c) and Monkey O (d). The correlation over the superficial electrodes is plotted in red, and the correlation over the deep electrodes is plotted in blue.

This pattern of deeper electrodes having more power than surface electrodes around the 10-20 Hz (low beta) region is also true of the post-stimulus period and is even more pronounced (Fig. 8a & 8b). Again in Monkey O, the pattern of deeper electrodes having more power than surface electrodes continues slightly past its peak frequency (Fig. 8b).

**Figure 8-.**
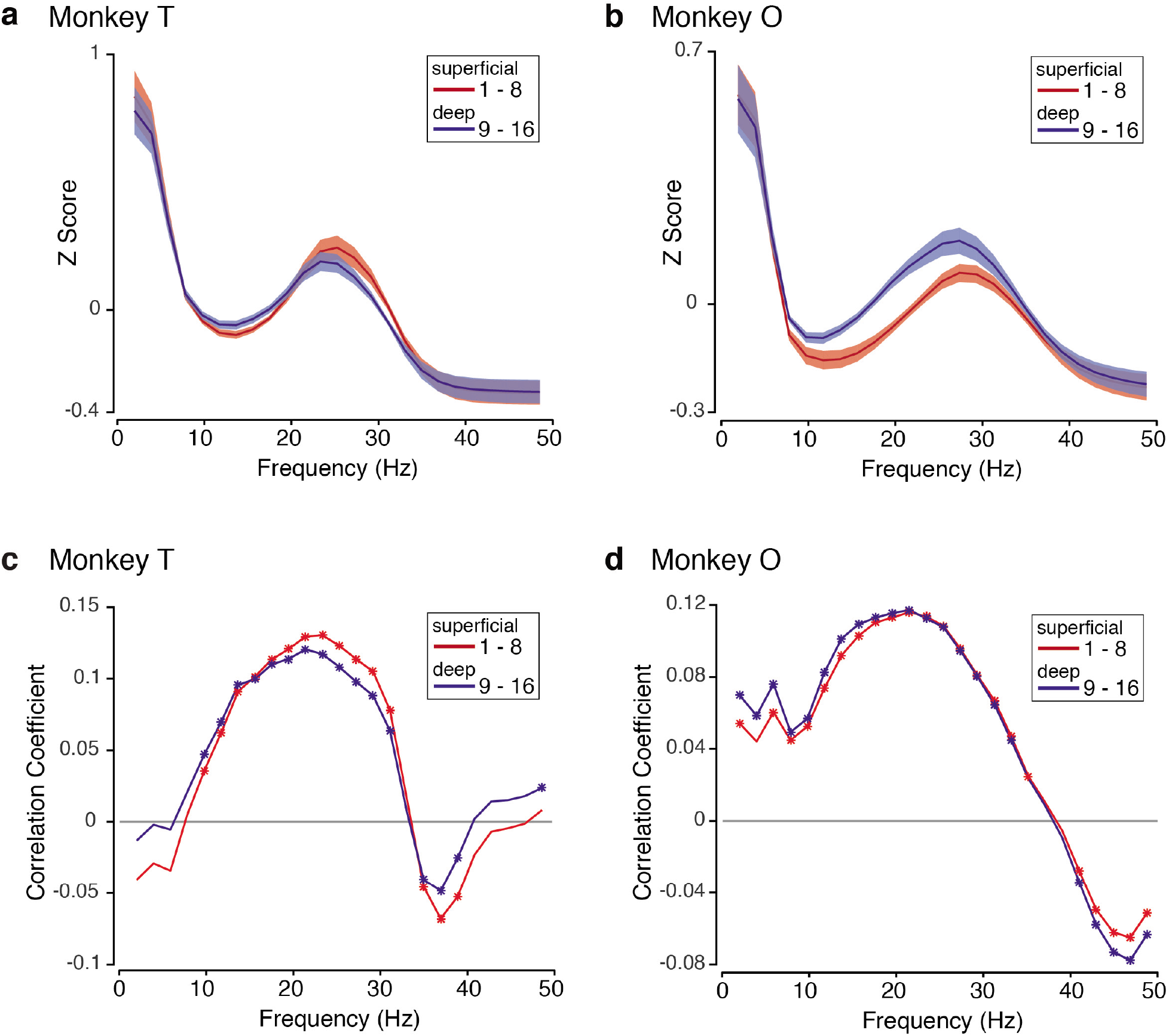
Post-stimulus BBA by depth. **a,b**: Normalized post-stimulus power spectra grouped into two electrode groupings and averaged over all trials, all electrodes within that group, and all sessions for Monkey T (a) and Monkey O (b). The power spectra have been normalized, and their Z Scores are plotted against frequency (Hertz). The average over the superficial electrodes is plotted in red, and the average over the deep electrodes is plotted in blue. Standard error over sessions is shaded. **c,d**: Depth dependent correlation between normalized post-stimulus power spectra with RT as a function of frequency for Monkey T (c) and Monkey O (d). The correlation over the superficial electrodes is plotted in red, and the correlation over the deep electrodes is plotted in blue.

### Correlation between BBA and RT does not vary significantly by depth

To examine whether BBA from certain cortical layers was more strongly tied with RTs, we performed the correlation with RT over two depth groups: superficial (electrodes 1:8) and deep (electrodes 9:16). For both pre-stimulus (Fig. 7c & 7d) and post-stimulus (Fig. 8c & 8d) BBA, the correlations for each group of electrodes produced the same shape as the correlation over all electrodes shown previously. The correlations for the superficial and deep electrodes are essentially the same, i.e. the correlation for one depth group is not significantly greater in magnitude than that of the other.

## DISCUSSION

The motivation for our study was to further understand the behavioral relevance of BBA and how it is organized as a function of cortical depth. In a perceptual decision-making task, we found that BBA was robustly present during the pre-stimulus and post-stimulus periods and was related to the behavioral RT. During the pre-stimulus period, low beta frequencies (∼15 to 20 Hz) were positively correlated with RT, while high beta frequencies (∼25 to 30 Hz) were negatively correlated. Through simulation, we found that the observed frequency-dependent correlation corresponds to a negative relationship between RT and the component frequencies of pre-stimulus BBA. During the post-stimulus period, all frequencies of BBA (∼15-30 Hz) were positively correlated to RT. We also found that deeper electrodes had higher power in the low beta frequencies (∼15 to 20 Hz) than superficial electrodes for both the pre- and post-stimulus periods.

### “Maintenance of current state” and “attentional” hypotheses help explain BBA in PMd

The nuanced relationship we discovered between BBA and RT is relevant for the ongoing discussion regarding the role of BBA. Currently, three main hypotheses exist, and each hypothesis has corresponding expected relationships between BBA and RT.

The postural hypothesis posits that BBA is a result of the maintained holding of a hand position and has no relationship to eventual behavior. For our experiment, one would predict no relationship between BBA and RT (Baker et al., 1999; Kristeva et al., 2007) – a hypothesis inconsistent with our findings that both pre-stimulus and post-stimulus BBA were related to RT.

Correlations between BBA and RT during the pre-stimulus period support both of the two remaining hypotheses. The maintenance hypothesis asserts that BBA represents a willingness to maintain the current state of either rest or movement. In this hypothesis, greater levels of BBA reflect the “desire” to maintain the hold position, which would result in slower movement and an increase in RT (Gilbertson et al., 2005; Pogosyan et al., 2009; Engel and Fries, 2010). Our finding of a positive correlation between BBA and RT for low beta frequencies is consistent with the maintenance hypothesis. The attentional hypothesis, which suggests that greater BBA reflects more attentional engagement with the task, would suggest a negative correlation between BBA and RT (Bouyer et al., 1987; Murthy and Fetz, 1992; Zhang et al., 2008; Saleh et al., 2010). The negative correlation between BBA and RT for high beta frequencies supports the attentional hypothesis.

During the post-stimulus period, we found that BBA was positively correlated with RT for both low and high frequencies, which supports the maintenance hypothesis. During this period, it appears that BBA of any frequency (low or high) reflects more willingness to maintain the current state of being.

This constellation of results suggest that the beta band is not a monolithic signal and consists of activity in at least two frequency sub-bands that dynamically emerge in different task epochs, perhaps reflecting distinct behavioral demands placed on the animal (Buschman et al., 2012; Spitzer and Haegens, 2017). We expand on this theme in the next section.

### BBA is better understood when split into two frequency bands

By examining the correlation at each frequency, rather than averaging over the whole beta frequency band, we found that BBA is better understood as being composed of at least two frequency sub-bands: low beta (∼13 to 20 Hz) and high beta (∼25 to 30 Hz).

Our nuanced view of BBA has some precedent in literature, with human EEG and rat studies referring to a beta1 band (∼ 15 Hz) and a beta2 band (∼ 25 Hz) (Haenschel et al., 2000; Kramer et al., 2008; Kopell et al., 2011; Cannon et al., 2014). In monkeys, Kilavik and collaborators examined motor cortical BBA during a visual multiple delay reaching task and suggested a similar separation (Kilavik et al., 2012). They posited that low beta frequencies were the result of widespread networks involved in top-down (conscious) processing and expectation of movement-related visual information, while higher beta frequencies emerged from bottom-up visual information processing and movement preparation (Kilavik et al., 2012).

The pre-stimulus period of our task incorporates the behavioral components identified by Kilavik and collaborators for both types of BBA – the monkey is expecting the visual checkerboard stimulus, is viewing relevant reach targets, and is preparing for one of two arm movements. We take the stance that the frequency composition of the pre-stimulus period reflects these different processes in the decision-making task. Therefore, it is not unreasonable that we see both low and high beta frequencies and positive and negative correlations between BBA and RT.

As the task progresses, the visual checkerboard (a bottom-up visual stimulus) appears. We speculate that the appearance of the checkerboard triggers a cognitive process that involves deliberation on the visual stimulus and likely movement preparation for the arm movement to report the decision. In the framework proposed by Kilavik and collaborators, such processes should induce activity in multiple beta frequencies, which is consistent with the broader frequency range of BBA we see in the post stimulus period. It remains to be understood why increased beta of any frequency during this period is associated with slower RTs.

### Beyond the LFP

Our study has focused on BBA in the LFP and behavior. We chose to analyze the LFP because it provides a population level, spatially averaged description of neural activity. We anticipate similar effects in spiking neurons, and preliminary analysis of our spike trains suggested BBA in many neurons and multi-units. However, analysis of single-neuron spike trains is often difficult because of the mixture of both poisson and non-poisson variability in these spike trains. Typical noise-reduction steps, such as convolution of spike trains with various filters, end up low pass filtering spike trains, which would lead to severe attenuation of signals at beta frequencies and the overemphasis of slower dynamics. We take the view these spikes are emerging from a dynamical system with activity at multiple time scales and that there is a need for collectively understanding both slow and fast dynamics in spiking activity. Single-trial analysis methods that use recurrent neural networks would facilitate such analyses (Pandarinath et al., 2017b).

### Greater low frequency beta in deeper electrodes is consistent with hypotheses about the generation of BBA

We found that electrodes placed deeper in the cortex, whose position approximately corresponds to layer V, have higher power in the low beta range (∼15 to 20 Hz) than superficially placed electrodes during both the pre- and post-stimulus periods. The power and depth relation differed across our two monkeys for high beta frequencies (∼25 to 30 Hz). The difference between monkeys for the power and depth relation in higher frequencies could arise due to variations in recording locations across animals or could be endogenous to the individual. This possibility would need to be studied with a variety of experiments and a larger test population.

Two main hypotheses exist regarding the generation of BBA: it is either generated locally, perhaps in layer V of motor cortex, or it is generated distally and transmitted from elsewhere (Khanna and Carmena, 2015; Spitzer and Haegens, 2017). Our finding of greater power in low beta frequencies for deeper electrodes is consistent with both predominant hypotheses; greater power could either indicate the BBA being generated in that layer (local hypothesis), or it could indicate that the distally generated BBA is projected into that layer (distal hypothesis).

Few studies have examined relationships between BBA and cortical depth. One study examined synchronization of BBA at various depths in the inferior temporal cortex during the passive repetition of visual stimuli (Kaliukhovich and Vogels, 2012). However, the passive nature of the task meant that they could not relate BBA to behavior. A recent laminar study of LFPs power in frontal cortex, including PMd, found greater power for low frequencies of BBA in deeper cortical layers (Bastos et al., 2018) – a result consistent with our observations here.

Even though few studies focus on how BBA changes as a function of cortical depth, many have hypothesized about its origin and built computational model s(Lee et al., 2013; Cannon et al., 2014). Despite these studies advancing our understanding of the biophysical basis of BBA, we still lack clarity about its underlying generators, because these modeling studies focus on results from in-vitro experiments in sensory cortices, with only one study focusing on the motor areas. Our study provides some of the first descriptions of BBA in premotor cortical areas in monkeys performing demanding cognitive tasks that also involve the somatomotor system. We anticipate that our data showing greater power in the lower frequencies of BBA will help constrain computational models of BBA. Studies involving laminar recordings in other BBA associated structures are needed to build the next generation of computational models of BBA. Ideally, these future studies would include decision-making, instructed delay, and somatosensory perturbation tasks that engage the different processes that are postulated to be associated with beta band activity.

## AUTHOR CONTRIBUTIONS

CC and KVS designed the experiments. CC performed the experiments, trained animals, and performed neurophysiological recordings. IEB and CC performed analyses together. IEB, CC, and KVS all wrote the paper. KVS was involved in all aspects of the manuscript.

## ACKNOWLEDGEMENTS

IEB was supported by the National Science Foundation Graduate Research Fellowship under Grant No. DGE-1656518. CC was supported by a NIH/NINDS K99/R00 award K99NS092972 and supported by HHMI as a research specialist. KVS was supported by the following awards: NIH Director’s Pioneer Award 8DP1HD075623, Defense Advanced Research Projects Agency (DARPA) Biological Technology Office (BTO) ‘‘NeuroFAST’’ award W911NF-14-2-0013, the Simons Foundation Collaboration on the Global Brain awards 325380 and 543045, and the Howard Hughes Medical Institute.

